## Supplemental Figure for "Real-time resolution of short-read assembly graph using ONT long reads"

### Supplementary Figures

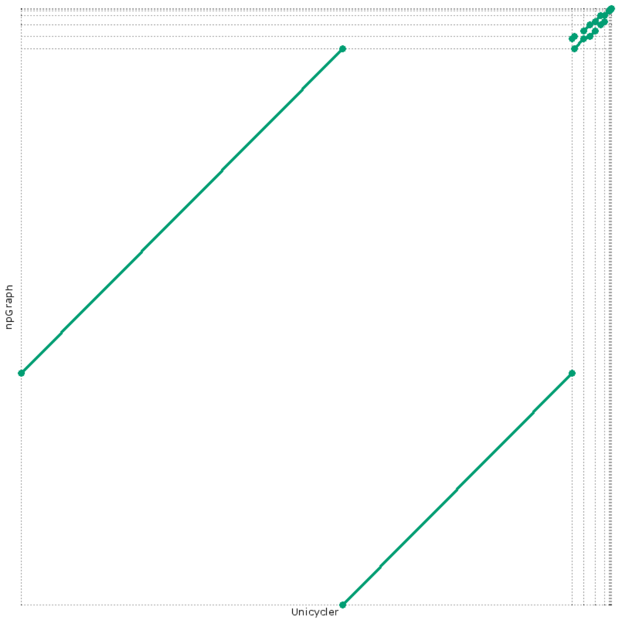


(a) *Citrobacter freundii* CAV1374

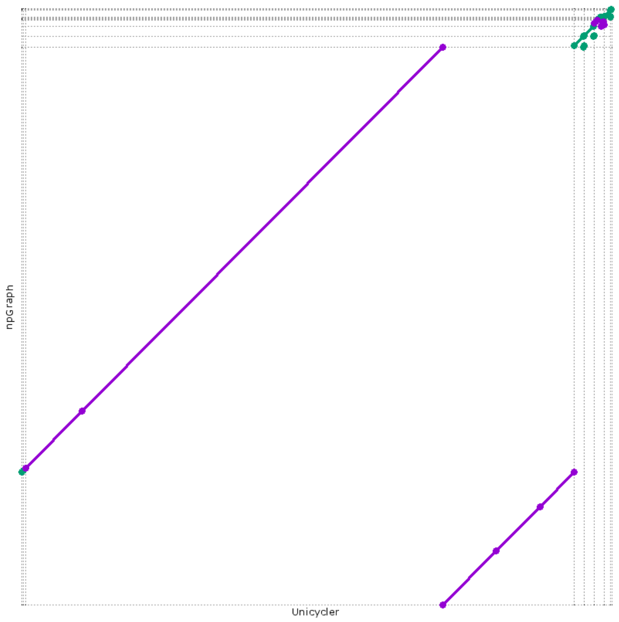


(b) *Klebsieall oxytoca* CAV1015

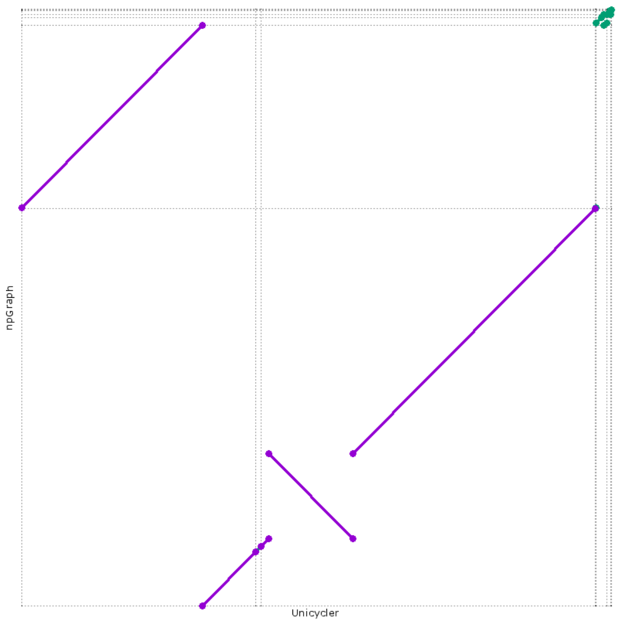


(c) *Enterobacter cloacae* CAV1411

Supplementary Figure 1: Dotplot generated by MUMmer for assembly results of `Unicycler` versus `npGraph`. Structural agreements between two methods were found in (a) *C.freundii* and (b) *K.oxytoca* assembly contigs. On the other hand, for (c) *E.cloacae* sample, there was a disagreement detected between 2 largest contigs given by two assembly algorithms. This case is investigated more thoroughly by using a reference from a same bacterial strain in Figure 2.

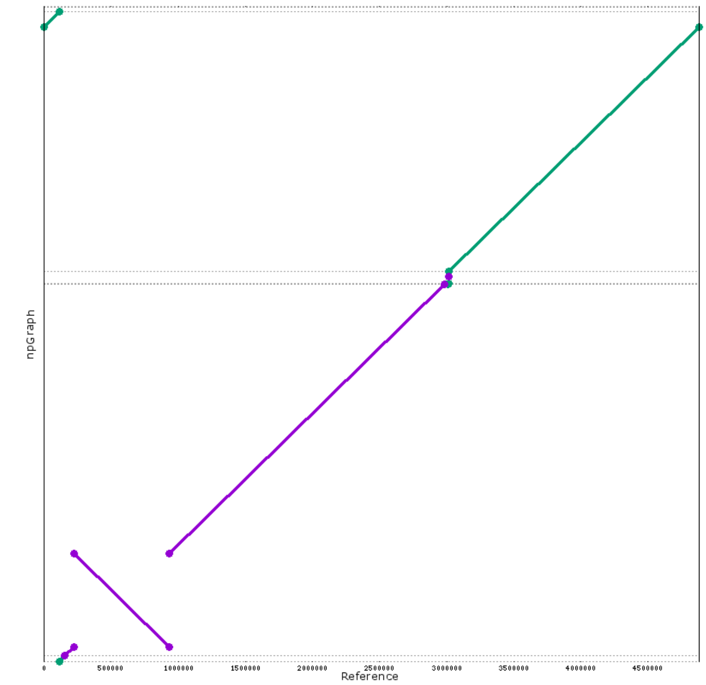


(a) *E. cloacae* `Unicycler` assembly versus reference genome

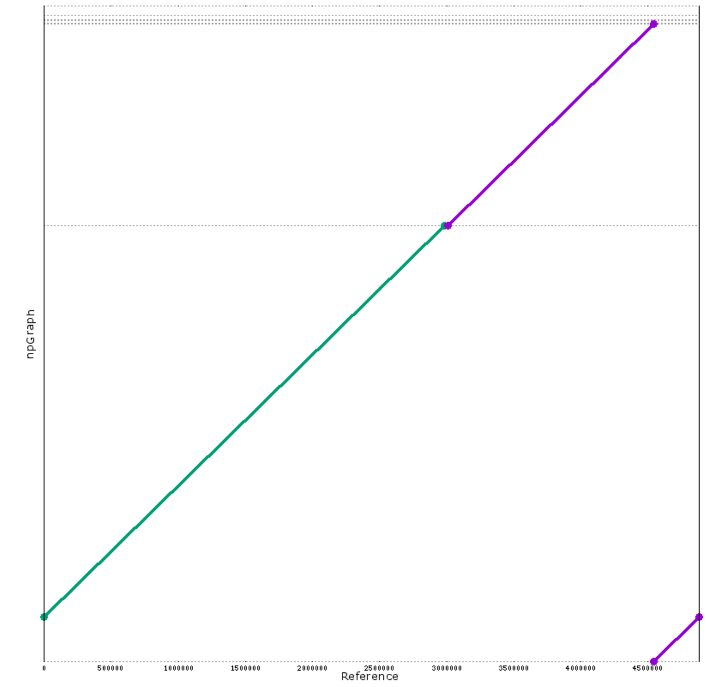


(b) *E. cloacae* `npGraph` assembly versus reference genome

Supplementary Figure 2: Alignments of an *Enterobacter cloacae* reference genome to assembly sequences generated by (a) `Unicycler` and (b) `npGraph`. While the former presents a structural variant, the latter is virtually an 1-to-1 mapping.
